# Immune response to vaccine candidates based on different types of nanoscaffolded RBD domain of the SARS-CoV-2 spike protein

**DOI:** 10.1101/2020.08.28.244269

**Authors:** Duško Lainšček, Tina Fink, Vida Forstnerič, Iva Hafner-Bratkovič, Sara Orehek, Žiga Strmšek, Mateja Manček-Keber, Peter Pečan, Hana Esih, Špela Malenšek, Jana Aupič, Petra Dekleva, Tjaša Plaper, Sara Vidmar, Lucija Kadunc, Mojca Benčina, Neža Omersa, Gregor Anderluh, Florence Pojer, Kelvin Lau, David Hacker, Bruno Correia, David Peterhoff, Ralf Wagner, Roman Jerala

## Abstract

Effective and safe vaccines against SARS-CoV-2 are highly desirable to prevent casualties and societal cost caused by Covid-19 pandemic. The receptor binding domain (RBD) of the surface-exposed spike protein of SARS-CoV-2 represents a suitable target for the induction of neutralizing antibodies upon vaccination. Small protein antigens typically induce weak immune response while particles measuring tens of nanometers are efficiently presented to B cell follicles and subsequently to follicular germinal center B cells in draining lymph nodes, where B cell proliferation and affinity maturation occurs. Here we prepared and analyzed the response to several DNA vaccines based on genetic fusions of RBD to four different scaffolding domains, namely to the foldon peptide, ferritin, lumazine synthase and β-annulus peptide, presenting from 6 to 60 copies of the RBD on each particle. Scaffolding strongly augmented the immune response with production of neutralizing antibodies and T cell response including cytotoxic lymphocytes in mice upon immunization with DNA plasmids. The most potent response was observed for the 24-residue β-annulus peptide scaffold that forms large soluble assemblies, that has the advantage of low immunogenicity in comparison to larger scaffolds. Our results support the advancement of this vaccine platform towards clinical trials.

## Introduction

Covid-19 is a pandemic viral disease caused by SARS-CoV-2 that emerged in 2019 and infected >23 millions of people across the world while the number of casualties is approaching 1 million ^1–3^ Since we currently lack an effective treatment of the disease and containment of the virus without imposing high cost for the society, vaccination seems to be the best hope to stop the waves of infection that continue to spread throughout the world. An effective vaccine should trigger formation of a protective humoral and cell mediated immune response against the viral components that will either inhibit viral entry and replication or kill virus-infected cells. Different vaccination platforms for the presentation of viral components have been used, including inactivated or attenuated viral particles, purified proteins, mRNA, plasmid DNA, nonreplicating viruses; each of them with particular features^4–9^. The advantages of DNA plasmid delivery including the speed of adaptation to new targets, cost effective production, stability at ambient temperature without the need for a cold chain make it a potentially attractive vaccination platform, although no DNA plasmid vaccine has been approved for humans so far^10,11^. Antibodies triggered by a vaccine should preferentially be focused to the domains and epitopes that can prevent viral recognition of the receptor, block viral fusion with cell membrane or interfere with viral replication in other ways. The majority of vaccines in current clinical trials are based on trimeric full length spike protein or its stabilized devivatives^5,9,12–18^, through which the virus attaches to the host cell receptor ACE2^19^. In this case antibodies against diverse surface exposed epitopes of the Spike protein are generated, where some of them may not prevent recognition of the ACE2 receptor or fusion and may even facilitate viral entry through an antibody dependent enhancement mechanism (ADE), as suggested before for SARS CoV and MERS CoV, as a highly undesirable property of a vaccine^20–22^. Therefore focusing immune response to the receptor binding domain (RBD) of the Spike protein may increase the probability of inducing neutralizing antibodies. Tertiary structure of the RBD in the complex with the ACE2 receptor has been determined^23,24^. Masking the receptor binding domain by the antibodies can prevent viral binding to the receptor and subsequent infection of target cells. Indeed it has been shown that RBD induces formation of neutralizing antibodies,^9,15^ similar as demonstrated for the related pathogenic coronaviruses^25,26^ and monoclonal antibodies targeting RBD have demonstrated effectiveness in preclinical studies^27,28^.

The ability of viral proteins or its domains to induce formation of antibodies depends on the structure and size of antigen. Viral surface proteins are typically presented to the immune system of the host in the form or nanoparticles tens of nm in diameter that present tens of copies of viral proteins. It has been shown that cells respond better to particles that present multiple copies of the antigen in comparison to the monomeric proteins, due to clustering of B cell receptors on B-lymphocytes, increased avidity of multimeric proteins and facilitated entry of particles above 20 nm to the lymph nodes^29–32^ Affinity maturation, class switching and memory cell formation takes place in germinal centers inside lymph node follicles^33^, therefore transport of protein antigens to lymph nodes is preferable in comparison to the insoluble protein antigen aggregates. It has been shown that nanoparticles above 50 nm are retained longer inside lymph node follicles and presented on the dendrites of follicular dendritic cells^34^. Presentation of the target antigen domain in the multimeric form may be accomplished by chemical conjugation of a viral protein domain to the nanoparticle or by a genetic fusion to the scaffold-forming polypeptide domain. Examples of the attachment of immunogenic domains to the virus-like capsule are capsid proteins of bacterial, plant or animal viruses such as Qβ, HPV, JCV, HBcAg, cowpea chlorotic mottle virus capsids^35^ and many others, nonviral protein compartments such as ferritin, lumazine synthase and encapsulin^31,36-39^ and *de novo* designed protein or DNA cages^40–46^.

Comparison of different scaffolding platforms could unravel which of them triggers the strongest response. An important issue of this strategy is that antibodies or cellular response may also be targeted against the scaffolding domain or a delivery vector, which could impair the efficiency of subsequent immunizations with the same vaccine type eliminate cells that produce the components of the scaffold proteins^47,48^. Implementation of the smallest scaffolding domain could therefore represent an advantage. Scaffolds used so far that form nanoparticles typically comprise 100-200 amino acid residues (e.g. ferritin, Qβ, lumazine synthase). On the other hand fusion to peptide tags with high aggregation propensity has also been used for vaccines^49–51^. In this case the amyloid or helical assembly-promoting peptide tag induced formation of large fibers that were decorated with peptide epitopes^52–54^. For the induction of most efficient antibody response it is important that the antigens readily reach the lymph nodes where affinity maturation of the adaptive immune response takes place^35^. Particles with sizes below ~500 nm can traffic through the lymphatic system into the germinal centers and can be taken up by antigen presenting cells such as dendritic cells^55^, therefore large insoluble aggregates are likely suboptimal.

Here we compared several strategies to present the RBD of the spike protein of SARS-CoV-2 on nanoscale particles, with stoichiometry in the range of 1 to >60 copies. Those assemblies were genetically encoded and implemented in animal studies as a DNA vaccine encoding fusion proteins for secretion from mammalian cells. We show that RBD fused to the scaffolding domains strongly increased the titer of RBD-specific antibodies that also recognized the Spike protein, neutralized interaction between the Spike and ACE2 receptor and provided protection in a surrogate infection assay. The proportion of the amino acid residues of the scaffold in those constructs ranged from as small as 8% to ~50% and those representing smallest fraction generated very low level of scaffold-directed antibodies even in 1 or 2 booster applications. Interestingly the response was best with fusion to a β-annulus peptide (RBD-bann) that had the propensity to generate large soluble oligomers rather than for other large structurally well defined scaffolds. Titers of antibodies against RBD after single immunization was most potent for larger RBD assemblies (RBD-bann and RBD-AaLS) and for all scaffolds substantially larger than for the monomeric RBD. This DNA vaccine also induced cytotoxic lymphocytes and γIFN producing lymphocytes. Our results warrant the advance of this type of vaccines into clinical trials. This type of nanoscaffolded assemblies could be implemented in either DNA, mRNA, viral or isolated protein-based platforms or combinations thereof in well considered prime-boost regimens.

## Methods

### Modelling of designed RBD-scaffolded protein cages

Molecular model structures of designed nanovaccines were prepared by performing homology modelling with Modeller (version 9.23)^56^. Structures PDB ID: 6VW and PDBID: 6VSB were used as templates for the receptor binding domain (RBD). For the RBD-ferritin construct, composed of 24 domains, PDB ID: 3EGM was used as template for the ferritin cage. RBD-foldon-RBD was modelled as a trimer based on the template PDB ID: 1RFO. For RBD-bann, homology modelling was used to first construct a model of the trimeric subunit based on the structure of the tomato bushy stunt virus (TBSV) (PDB ID: 2TBV). Twenty trimeric subunits were then assembled into a RBD-decorated protein cage by employing icosahedral symmetry characteristic for TBSV, resulting in a 60-mer with a diameter of approximately 26 nm. Model of the design RBD-AaLS, composed from 60 subunits, was built by employing PDBID: 1HQK as template for the lumazine synthase scaffold.

### Preparation of DNA constructs

DNA constructs were prepared using conventional methods based either on synthetic DNA (Twist, IDT) or purchased commercially (Genscript) or clones from plasmids with viral proteins generously provided by Prof. Nevan Krogan, UCSF^57^. The construct encoding prefusion ectodomain of the SARS-CoV-2 Spike protein was a generous gift from Prof. Jason McLellan, University of Texas, Austin^23^. The RBD domain of Spike (residues R319-S591) was codon-optimized for *H. sapiens* and synthesized (Genscript) into pcDNA3.1 (+) with a human pregnancy specific glycoprotein 1 signal peptide at the N-terminus and a 3C-protease cleavage site followed by a His-tag at the C-terminus. ACE-2 (hsACE-2, residues S19 to D615) was codon optimized for *H. sapiens* and cloned into the pTwist_CMV_BetaGlobin_WPRE_Neo (Twist Biosciences). It is preceded with a human pregnancy specific glycoprotein 1 signal peptide, and C-terminally tagged with a 3C protease cleavage site, twin-strep tag and 10x His tag.RBD domain of the Spike protein encompassing residues P330 to K521 was fused with polypeptide scaffolds and inserted in pcDNA3.1 (+) vector with a CD45 signal peptide at the N-terminus. β -annulus scaffold peptide from the tomato bushy stunt virus^58^, RBD-bann, chimeric fusion of Bullfrog *(Rana catesbeiana)* and *Helicobacter pylori* ferritin^59^, (RBD-ferritin), lumazine synthase from *Aequifex aeolicus^60^,* (RBD-AaLS) and foldon from T4 bacteriophage fibiritin, (RBD-foldon-RBD), were codon optimized for expression in mammalian cells. Non-tagged versions of RBD-scaffold constructs were prepared in parallel for immunization purposes.

### Cell culture

The human embryonic kidney (HEK) 293, HEK293T (ATCC) cell line and mouse NIH-3T3 cell (ATCC) line was cultured in DMEM (Invitrogen) supplemented with 10% FBS (BioWhittaker) at 37 °C in a 5% CO_2_ environment. ExpiCHO cells (Thermo Fisher) were cultured in ProCHO5 medium (Lonza) and incubated with agitation at 31 °C and 4.5% CO2. HEK293E cells were cultured in EX-CELL 293 serum-free medium (Sigma) and incubated with agitation at 37°C and 4.5% CO_2_. Expi293F cells (Thermo Fisher) were cultivated in Expi293™ Expression Medium (Thermo Fisher) at 37°C and 8% CO _2_ on an orbital shaking platform with shaking speed based on shaking diameter and flasks’ size.

### Recombinant viral proteins

The prefusion ectodomain of the SARS-CoV-2 Spike protein was transiently transfected into suspension-adapted ExpiCHO cells (Thermo Fisher) with PEI MAX (Polysciences) in ProCHO5 medium (Lonza). After 1 h, dimethyl sulfoxide (DMSO; AppliChem) was added to 2% (v/v). Incubation with agitation was performed at 31°C and 4.5% CO_2_ for 5 days. The clarified supernatant was purified via a Strep-Tactin column (IBA Lifesciences) and dialyzed into PBS buffer. Average yield for Spike was 15 mg/L culture.

The RBD domain of Spike (RBD, residues R319-S591) was transiently transfected into suspension-adapted ExpiCHO cells (Thermo Fisher) with PEI MAX (Polysciences) in ProCHO5 medium (Lonza). After 1 h, dimethyl sulfoxide (DMSO; AppliChem) was added to 2% (v/v). Incubation with agitation was performed at 31°C and 4.5% CO2 for 6 days. The clarified supernatant was purified in two steps; via a Fastback Ni2+ Advance resin (Protein Ark) followed by Superdex 200 16/600 column (GE Healthcare) and finally dialyzed into PBS buffer. Average yield for RBD-His was around 25 mg/L culture.

ACE-2 protein (hsACE-2, residues S19 to D615) was transiently transfected into HEK293E cells (Thermo Fisher) with PEI MAX (Polysciences) in EX-CELL 293 serum-free medium (Sigma), supplemented with 3.75 mM valproic acid. Incubation with agitation was performed at 37°C and 4.5% CO_2_ for 8 days. The clarified supernatant was purified in three steps; via a Fastback Ni2+ Advance resin (Protein Ark), a Strep-Tactin XT column (IBA Lifesciences) followed by Superdex 200 16/600 column (GE Healthcare) and finally dialyzed into PBS buffer. Average yield for hsACE-2 was 12 mg/L culture.

### Production and characterization of RBD variants

Expi293F (Thermo Fisher) cells were transiently transfected with ExpiFectamine™ 293 Transfection Kit (Thermo Fisher) according to manufacturer’s recommendations, and grown in Expi293™ expression medium (Thermo Fisher). Incubation with agitation was performed at 37°C and 8% CO2 for 5 days. For proteins containing Histag, cell broth was collected after 5 days and clarified by centrifugation, followed by overnight dialysis for against NiNTA A buffer (50 mM Tris, 150 mM NaCl, 10 mM imidazole, pH 8) with dialysis tubes, 6-8 kDa cutoff (Spectrum Laboratories, USA). Samples were purified using NiNTA resin (Goldbio, USA) and eluted with 250 mM imidazole in NiNTA A buffer. For proteins containing strep-tag, cell broth was also collected after 5 days and clarified by centrifugation, followed by purification, using Strep-trap (Cytiva, USA), according to manufacturer’s recommendations. After affinity isolation all samples were dialyzed against 20 mM Tris,150 mM NaCl, pH 7.5.

### Dynamic light scattering (DLS)

Particle size was measured on a ZetasizerNano (Malvern, UK) at 20 °C using an angle of 173° and 633-nm laser. Measured data was recorded and analyzed with the software package, provided by the manufacturer.

### Size exclusion chromatography coupled to light scattering

SEC-MALS analysis was performed with Superdex 200 increase 10/300 column on HPLC system (Waters, USA), coupled to a UV detector (Waters, USA), a Dawn8+ multiple-angle light scattering detector (Wyatt, USA) and a RI500 refractive index detector (Shodex, Japan). Samples were filtered through 0.2 μm centrifuge filters (), and injected onto previously equilibrated column (20 mM Tris, 150 mM NaCl, pH 7.5). Astra 7.0 (Wyatt, USA) was used for data analysis.

### ELISA assay

Mammalian cell line HEK293 was transfected with plasmids for vaccination coding fusion proteins of the RBD to the scaffolding domains in order to produce in cells the protein immunogen. HEK293 cells was cultured in DMEM (Invitrogen) supplemented with 10% FBS at 37 °C and 5% CO2 environment. 24h prior transfection cells were seeded at 2 x 105 cells/ml. At a confluence of 40-70 %, cells were transfected with a mixture of DNA and PEI. For every 1 μg of DNA transfected, 6 μL PEI at stock concentration of 0,324 mg/ml, pH 7,5 was diluted in 150 mM NaCl. The mixtures of DNA and PEI were incubated at room temperature for 15 min and added to the cell medium. Supernatants of HEK293 were harvested 3 days post transfection and analyzed with ELISA using anti-RBD antibodies to determine the expression profile of proteins (Figure 2). For ELISA assay high binding half-well plates (Greiner) were coated with antibody SARS-CoV-2 (2019-nCoV) Spike Antibody (Sinobiological; 40150-R007; diluted 1:500) and incubated overnight at 4°C. Next, plates were washed with 1xPBS + 0,05 % Tween20 and blocked for 1h at RT with 100 μl of ELISA/ELISPOT diluent solution (eBioscience). After washing samples were incubated for 2h at room temperature with 1X ELISPOT Diluent. Then followed the steps of washing, detection Ab Tetra His Antibody, BSA free (Qiagen; 34670; diluted 1:2000; 1h room temperature) and washing. After the addition of the substrate (TMB solution) the reaction was stopped with 0,16 M sulfuric acid. Multiplate reader SineryMx (BioTek) was used to measure absorbance. Absorbance at 620 nm was used for correction and was subtracted from the absorbance at 450 nm.

### Mouse immunization studies

To test the immunogenicity of DNA vaccines female 8-10 weeks BALB/c OlaHsd mice (Envigo, Italy) were used for immunization protocols. All animal experiments were performed according to the directives of the EU 2010/63 and were approved by the Administration of the Republic of Slovenia for Food Safety, Veterinary and Plant Protection of the Ministry of Agriculture, Forestry and Foods, Republic of Slovenia (Permit Number U34401-8/2020/9, U34401-8/2020/15, U34401-12/2020/6).

Laboratory animals were housed in IVC cages (Techniplast), fed standard chow (Mucedola) and tap water was provided ad libitum. Mice were maintained in 12-12 hours dark-light cycle. All animals, used in the study, were healthy; accompanied with health certificate from the animal vendor.

Immunization was carried out under general inhalation (1,8% MAK isoflurane anesthesia (Harvard Apparatus)). Immunization protocol was based, if not stated otherwise, on prime vaccination and two boosts with 2-week interval between vaccinations. Animals were vaccinated with plasmid DNA (RBD, RBD-scaffold, scaffold vectors, empty pcDNA3.1 (+) vector), either naked DNA or coupled with jetPEI-in vivo transfection reagent (Polyplus Transfection) with the N/P ratio 12. Each animal received a total of 20 μg of designated plasmid DNA, combined with transfection reagent via intramuscular injection (total volume of 50 μl per animal).

In stated experiments, animals received naked plasmid DNA or isolated recombinant proteins (30 μg/animal of designated recombinant protein, coupled with/without alum adjuvant (40 μg/animal) Alhydrogel^®^ (Invivogen; vac-alu-250).

Vaccine was administered using 30G needle (Beckton Dickinson) into m.tibialis anterior after appropriate area preparation. One day before each boost, blood was drawn from lateral tail vain using Microvette 300 (Sarstedt). Two weeks after the second boost, the experiment was terminated. Final blood was taken and spleens were harvested from animals for further analysis. Mouse sera were prepared by centrifugation of blood samples 3000RPM/ 20min at 4°C. In mouse sera specific mouse antibodies were determined by ELISA in order to analyze the immunogenicity of designed RBD DNA vaccines. In different experiment switch immunization was performed, where prime and boost was carried out with RBD DNA, fused to two different scaffolds, e.g. prime was carried out with bann-RBD and for the boost RBD-foldon-RBD was used and vice versa.

### Analysis of immune response on mice

ELISA tests were performed to determine endpoint titer for designated specific antibodies. High binding half-well plates (Greiner) were used. According to desired total IgG determination, appropriate recombinant protein was used for ELISA plate coating. Recombinant proteins were coated in PBS buffer (Gibco) at the concentration of 1.2 mg/ml of protein per well. Coated plates were incubated overnight at 4°C. Next day plates were washed with PBS+0.05 % Tween20 (PBS-T) using ELISA plate washer (Tecan). Next, plates were blocked for 1h at RT with 100 μl of ELISA/ELISPOT diluent solution (eBioscience). Afterwards, plates were again washed. Then serial dilution of sera was added to plates, where each dilution presented a certain titer values. In the first row a 1:100 dilution was added, then a 3-fold dilution was performed with each row. For mouse sera ELISA diluent was used. Mouse sera were incubated at 4°C overnight. Next day, plates were washed and afterwards specific secondary antibodies (dilution 1:3000), coupled with HRP were added to wells. For total IgG determination, goat anti mouse IgG (H+L)-HRP antibodies (Jackson ImmunoResearch; 115-035-003) were used. To determine specific types of anti-RBD antibodies in mouse sera goat anti-mouse IgG1-HRP (Abcam; ab97240), goat anti-mouse IgG2a heavy chain-HRP (Abcam; ab97245), goat anti-mouse IgG2b heavy chain-HRP (Abcam; ab97250), goat anti-mouse IgG3 heavy chain-HRP (Abcam; ab97260), goat anti-mouse IgM mu chain-HRP (Abcam; ab97230), goat anti-mouse IgA alpha chain-HRP (Abcam; ab97235) were used. Plates were incubated for 1h at RT. Next, plates were washed. After the final wash, TMB substrate (Sigma) was added and the reaction was stopped with the addition of acid solution (3M H3PO4). Absorbance (A450 nm, A620 nm) was measured by Synergy Mx microtiter plate reader (Biotek). End point titer was determined as the dilution above the value of the cutoff. From absorbance data of control animals (vaccinated with empty vector pcDNA3.1(+)) the cutoff value was determined^62^. For confidence level of 95% value 2,335 was used as the standard deviation multiplier.

### VSVΔG* pseudotyped virus system and pseudoviral neutralization assay

Pseudovirus system based on the vesicular stomatitis virus^63^ was used to determine virus neutralization capacity of immunized mice sera, antigen specific cellular immunity and animal infection studies. Plasmids and VSVΔG*/G virus were kindly provided by Stefan Pölhmann. Pseudovirus preparation was described previously^19^. Briefly, HEK293T cells were seeded into 6-well plates (9*105/well) a day before transfection with pCG1-Spike/PEI mixture. The next day, cells were infected with VSVΔG*/G virus in serum-free medium for 1h, after which medium was removed and cells were washed with PBS before complete medium supplemented with anti-VSV-G antibody (8G5F11, Kerafast) was added to the cells. After 18h cell supernatant was centrifuged and cleared pseudovirus supernatant was aliquoted and stored at −80°C until use.

For neutralization assay, HEK293 were seeded (2.5*104/well) a day before transfection with plasmids encoding ACE2 (pCG1-ACE2, a kind gift of Stefan Pölhmann), pCMV3-C-Myc-TMPRSS2 and phRL-TK (encoding Renilla luciferase). Immunized mice sera were preincubated with spike pseudovirus for 30min before addition to the cells. The next day, medium was removed and cells were lysed in Passive lysis buffer (Biotium). Luciferin substrate (Xenogen) was used to detect Firefly luciferase activity as a measure of pseudovirus infection and coelenterazine H (Xenogen) to follow Renilla luciferase activity for determination of transfection efficiency and normalization.

### Neutralization assay based on inhibition of ACE2 Spike interaction

Recombinant human ACE2-Fc (Genscript) at concentration of 2 μg/ml in phosphate buffer saline (PBS) was adsorbed to wells of ELISA plates at 4°C overnight. Serum dilutions were prepared in 1% BSA/PBS-T and incubated with Spike protein at final concentration of 0.1 μg/ml for 1h at 37°C. After blocking in 1% BSA/PBS-T for 1 h at 37°C plates were washed in PBS-T and serum-Spike samples were added to wells and incubated for 2h at room temperature. After wash the plate was incubated with streptactin-HRP (1:10000) in 1% BSA/ PBS-T for 1h at room temperature. After the final wash, TMB substrate was added and the reaction was stopped by the addition of acid solution (3M H_3_PO_4_). Absorbance was measured by Synergy Mx microtiter plate reader (Biotek).

### T cell response on mouse splenocytes

To determine the presence of antigen-specific cytotoxic CD8a+ T cells, mice spleens from RBD DNA or RBD-scaffold DNA immunized animals were harvested. Single cell suspensions from spleens without enzymatic treatment were obtained using tissue dissociator gentleMACS™ Dissociator, according to the manufacturer’s instruction (Miltenyi Biotec). CD8^+^ T cell from spleen cell suspension were isolated using mouse CD8a^+^ T Cell Isolation Kit (Miltenyi Biotec; 130-104-075) according to the manufacturer’s instruction. Cells were isolated based on negative selection using LS columns, obtaining up to 10^8^ labelled cells. To determine RBD specific cytotoxicity, mouse NIH-3T3 cells were seeded into 24-well plates (1*10^5^/well); next day cells were transfected with pCG1-hACE2 (900 ng/well) and pCMV-TMPRSS2 (30 ng/well) plasmids. The following day, cells were infected with Spike pseudovirus with bioluminescent reporter. Next day isolated CD8a+T cells (1*10^5^/well) were added in RPMI1640 cell medium. After 24 hours, bioluminescence was determined using IVISIII (Perkin Elmer) after the addition of D-luciferin (500 μM), showing the state of RBD-specific cytotoxicity of CD8+T cells, isolated from RBD DNA or RBD-scaffold DNA vaccinated animals. Bioluminescence values are presented as Average Radiance (p/s/cm^2^/sr), which were determined using Living Image^®^ software. From Average Radiance values (ARV) the percentage of infected NIH-3T3 specific lysis was calculated using formula: % specific lysis=100*(spontaneous death ARV-test ARV)/(spontaneous death ARV-maximal killing ARV).

To determine cell specific response, mouse splenocytes were seeded into 24-well plates (1*10^6^/well). Cells were then stimulated with SARS-CoV-2 (Spike Glycoprotein) PEP-LIPS-Pool peptide pool (peptides&elephants GmbH; LB01792), consisting out of 316 peptides at final concentration of 10 μg/ml. Next day, mouse IFNγ was determined, using Mouse IFN gamma ELISPOT ELISA Ready-SET-Go!™ Kit; 15531137) according to manufacturer’s instructions.

### Surrogate assay of protection of viral infection by immunization

Balb/c mice were immunized with RBD-bann according to scheme, presented above. Two weeks after the boost, mice were transfected by 30 μl of plasmid mixture of jetPEI- in vivo and plasmid DNA (20 μg hACE2, 1 μg TMPRRS per animal) via intranasaly administration in inhalation anesthesia. Next day, mice were again, intranasal, infected with 70 μl of VSV-S pseudovirus. 24 hours later, mice received 150 mg/kg of body weight of D-luciferin (Xenogen) intraperitoneally and were *in vivo* imaged by using IVIS^®^ Lumina Series III (Perkin Elmer). Bioluminescence that depicted the state of pseudovirus infection was determined. Data were analyzed with Living Image^®^ 4.5.2 (Perkin Elmer).

## Results

### Design of RBD-presenting polypeptide nanoparticle scaffolds of different size and oligomerization state

Fusion of protein antigens to the self-assembling scaffold can generate a range of stoichiometries and sizes that may have a varying effect on the immune response. In case of SARS-CoV-2 vaccines trimeric S protein has been used in many constructs resulting in protein complexes of a molecular weight exceeding 400 kDa. Additionally a dimeric and trimeric version of the RBD domain demonstrated increased immunogenicity^25,15^ and VLPs based on the chemical crosslinking of RBD to the viral capsids that present multiple copies of RBD domains have been reported.^64^ Genetically encoded fusion between the protein antigen and a scaffold polypeptide enables the use of diverse vaccine platforms based on nucleic acids, similar as demonstrated by the ferritin-based vaccine against influenza, HIV-1 and RSV^38,65^. Aiming to explore the influence of the number of RBDs per particle and minimize the size of the scaffold in order to reduce the immune response against the scaffold, several scaffolding variants were engineered. We decided to compare the immune response against the RBD domain presented in 1, 6, 24, and ≥60 copies. Besides generating a range of antigen stoichiometries, the generated particles should present the antigen at their surface and preferably shield the scaffold from recognition by antibodies. This was achieved by fusing the RBD to the foldon (RBD-foldon-RBD), ferritin (RBD-ferritin), lumazine synthase (RBD-AaLs) and β-annulus peptide (RBD-bann). Those scaffolds range in size from 24 to 173 amino acid residues, which means that they represent between 7-46% of amino acid residues in the genetically engineered fusion constructs. The smallest stoichiometry of the assembly in our set, six, was achieved in the RBD-foldon-RBD construct, where the RBD domain was fused to both amino- and carboxy-termini of the trimerizing foldon domain^66^. Since the RDB is much larger than the foldon peptide it can, according to the molecular model (Fig. 1), more efficiently shield the foldon trimerization core in comparison to the conventionally used trimeric fusion at only one end. An additional potential advantage of our designs is to support trimerization of the RBD, which mimics the arrangement on the trimeric Spike. Higher stoichiometries of the assemblies can even more efficiently mask scaffold domains. Ferritin forms a 24-meric cage and has been frequently used for the presentation of polypeptide antigens as it can present 8 trimers on each particle^38,65,39^. However, the density of RBD domains at its surface is, according to the molecular model, lower than for the other designed assemblies (Fig. 1). Even larger number of RBD domains were presented on the icosahedral *Aequifex aeolicus* lumazine synthase (AaLs), which has also been used before for nanostructured vaccines^60,67^. A short peptide, comprising 24 amino acid residues based on the viral β-annulus protein from tomato bushy stunt virus has been shown to spontaneously form nanoparticles approximately 40 nm in diameter^58^. This peptide has been used before to attach different small cargo molecules, from DNA to peptides^68,69,70^. We hypothesized that this peptide could be fused to a large folded protein domain to generate formations of large soluble assemblies for vaccination. The molecular model was constructed according to the icosahedral packing of its parent viral capsid, which comprises 60 copies, however, for this construct, the experimental results suggested larger polydisperse soluble assemblies (see below).

**Figure 1:**
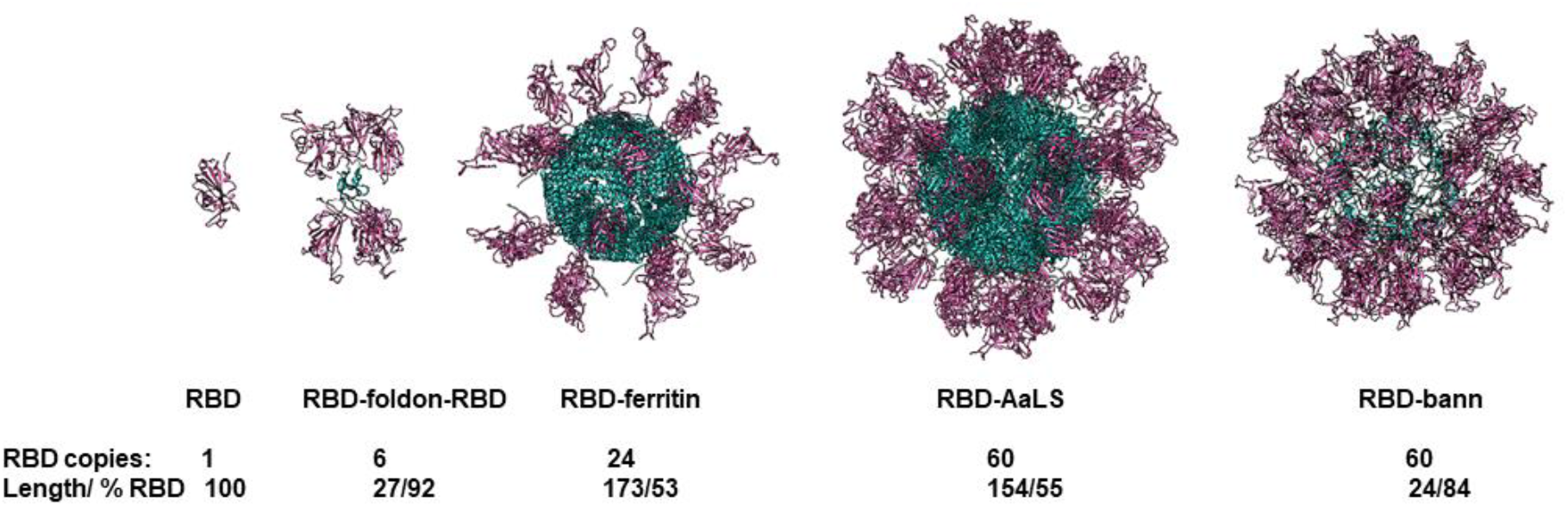
Molecular models of designed scaffolded RBD domains. RBD domains are shown in violet and scaffold core in blue. Number of RBD domains per particle, length of the scaffolding domain and fraction of amino acid residues the RBD in the assembly are written below each model.

Constructs for the genetically encoded scaffolded RBD were implemented as DNA vaccines and included a signal sequence for protein secretion as the cytoplasmic expression has low efficiency of immune response^71^. The analysis of a supernatant of the human cell line HEK293 and ExpiCHO transfected with appropriate plasmids revealed that all proteins were produced and secreted into the medium (Fig. 2a). Interestingly, determination of the size of purified RBD-bann fusion protein secreted from mammalian cells revealed that the size of the assemblies was larger as predicted, with particles around 500 nm (Fig. 2b), however, the size in the tissue milieu may be affected by the physiological conditions.

**Figure 2:**
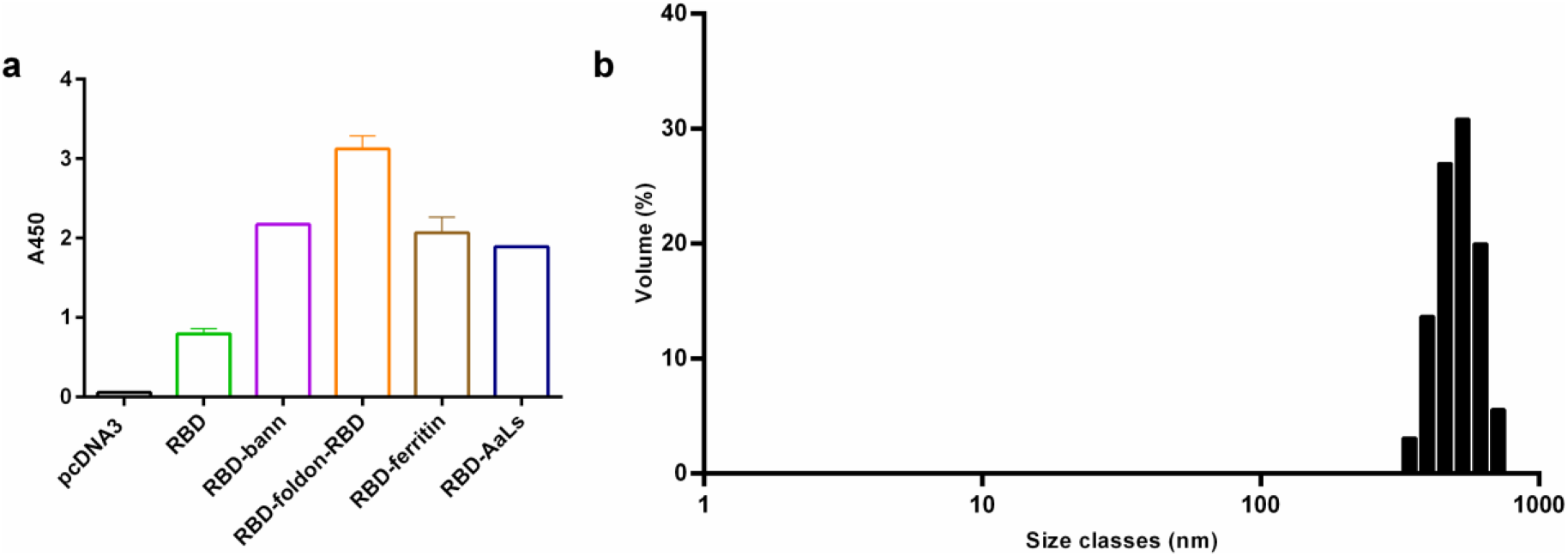
Secretion of RDB protein domains fused to different scaffolding proteins produced in plasmid-transfected mammalian cells and size analysis of the isolated RBD-bann protein. Supernatant of HEK293 cells transfected with indicated construct was harvested 3 days post transfection and the presence of differently scaffolded RBD domain variants was detected with anti RBD antibodies (A). Size analysis of the purified RBD-bann by DLS confirms the presence of particles around 500 nm (B).

### Immune response to DNA vaccines coding for the scaffolded RBDs

Purified endotoxin-free plasmids were used for immunization of mice. We followed the scheme of intramuscular priming followed by the subsequent booster immunizations after two weeks (Fig. 3a). Samples of blood were collected before the subsequent injection of plasmid DNA and two weeks after the final immunization, animals were sacrificed and used for the analysis of the immune response. Titer of antibodies two weeks after the priming immunization differed depending on the type of the scaffold and tested antigen. The weakest response of antibodies recognizing RBD was obtained against the monomeric RBD (Fig. 3b-d). The response was best for the RBD-bann and RBD-AaLs and weaker against the RBD-foldon-RBD and RBD-ferritin. However titers of antibodies against the Spike protein were similar for all scaffolded RBDs, two orders of magnitude better than against the monomeric RBD (Fig. 3e-g). After the first booster immunization titers were even more similar for the scaffolded RBD, where they increased from 95- for RBD-AaLs, 130- for RBD-foldon-RBD, 150- for RBD-ferritin to 180-fold for RBD-bann, all of them orders of magnitude higher than for the monomeric RBD. The third immunization only increased the titer for the monomeric RBD but not for others, demonstrating that the two immunizations should be sufficient to reach the plateau response. Analysis of the antibody classes (Fig.4) showed mainly IgG1, IgG2b and IgG3 with lower titers of IgM and IgA and no IgG2a, suggesting Th1-biased response, typically observed for the DNA vaccines^72^.

**Figure 3:**
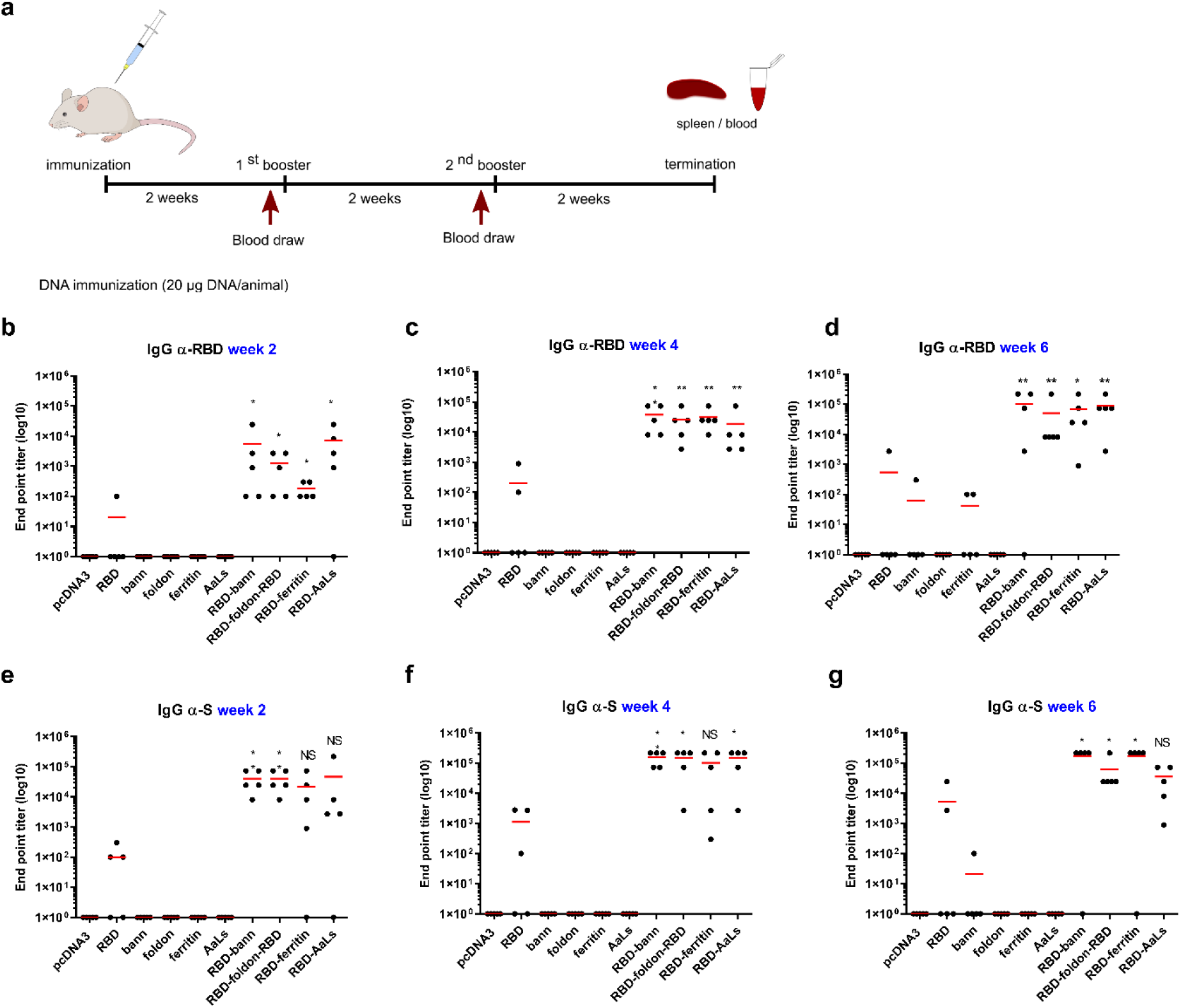
Titer of total IgG antibodies against the RBD and Spike protein for immunization with plasmids for different scaffolded RBDs and scaffold alone. Mice were immunized with different combination of RBD plasmid DNA, complexed with jetPEI-*in vivo* transfection reagent, according to immunization protocol (A). End point titer (EPT) for total IgG against RBD (B-D) and against Spike protein (E-G). Graphs represent mean of EPT of group of mice (n=5 per group). Each dot represents an individual animal. *P< 0.05; **P < 0.01. All P values are from Mann-Whitney test.

**Figure 4:**
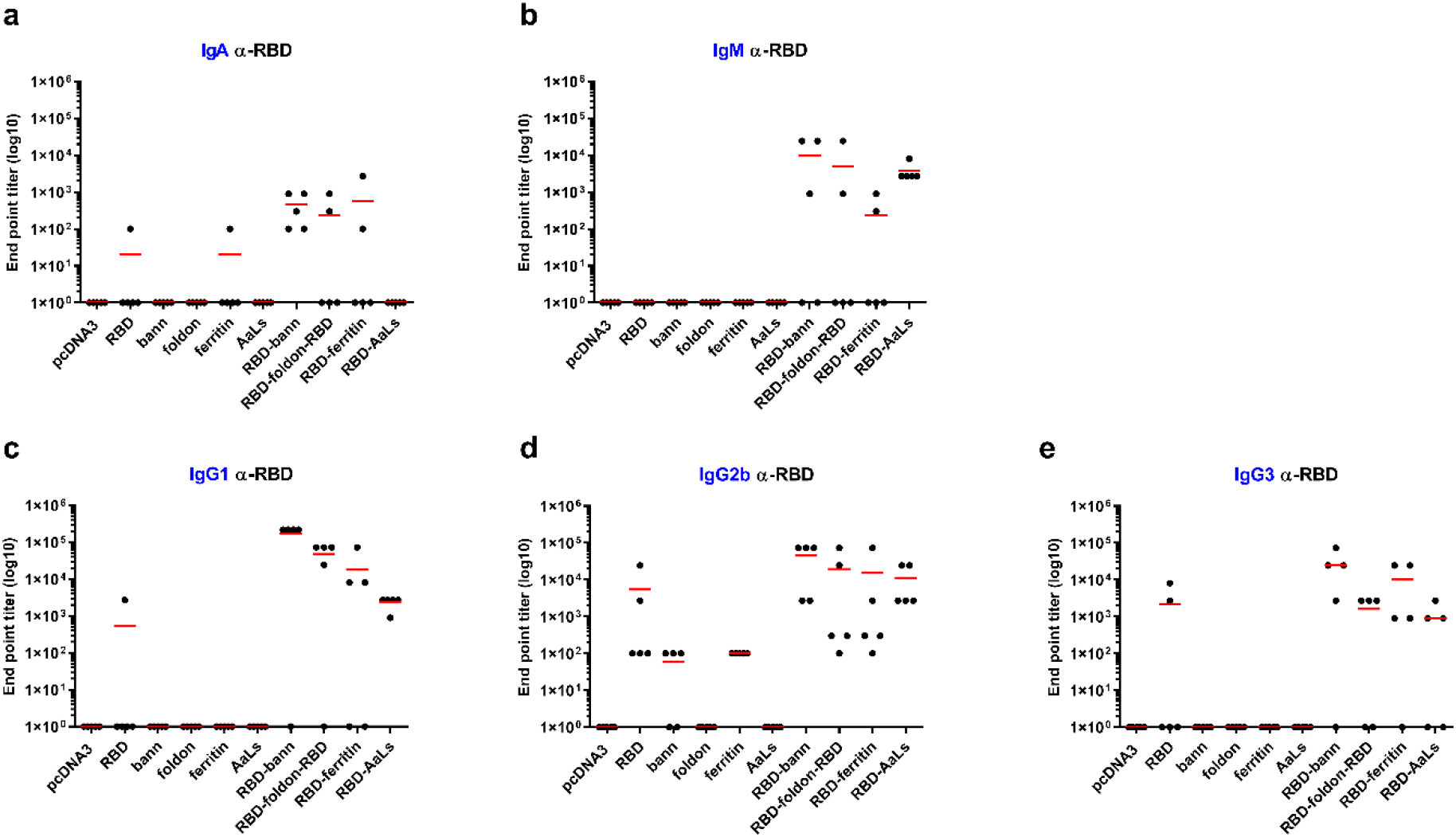
Analysis of different classes of antibodies against RBD for different scaffolded RBDs and immunization by the scaffold. Mice were immunized with different combination of RBD plasmid DNA. End point titers 6 weeks after the first immunization of IgA (A), IgM (B), IgG1 (C), IgG2b (D) and IgG3 (E) against RBD protein were determined by ELISA. Graphs represent mean of EPT of group of mice (n=5 per group). Each dot represents an individual animal.

While plasmids in combination with jetPEI transfection gent were used initially, further immunization experiments demonstrated that the use of naked plasmid DNA without added jetPEI did not decrease the antibody titer compared to immunization with complexed jetPEI-DNA mixture (Supplementary Figure 1).

### Neutralization of viral protein binding to the receptor by sera

Neutralization of the viral engagement of the ACE2 receptor by the Spike protein was investigated by a neutralization test based on the inhibition of interaction between Spike and ACE2, that has been shown before to correlate well with neutralization of viral infection^73^. Results showed best neutralization by the sera of mice immunized by RBD-bann vaccine with IC50 dilution of ~1:220 (Fig. 5a, Sup.Fig. 2). Viral neutralization was additionally tested using S-pseudotyped VSV viral assay (Fig. 5b). Similar as in the interaction neutralization assay, best neutralization was obtained for the RBD-bann, followed by RBD-foldon-RBD, RBD-ferritin and RBD-AaLs. In both assays protection was low for immunization by DNA coding for RBD monomer.

**Figure 5:**
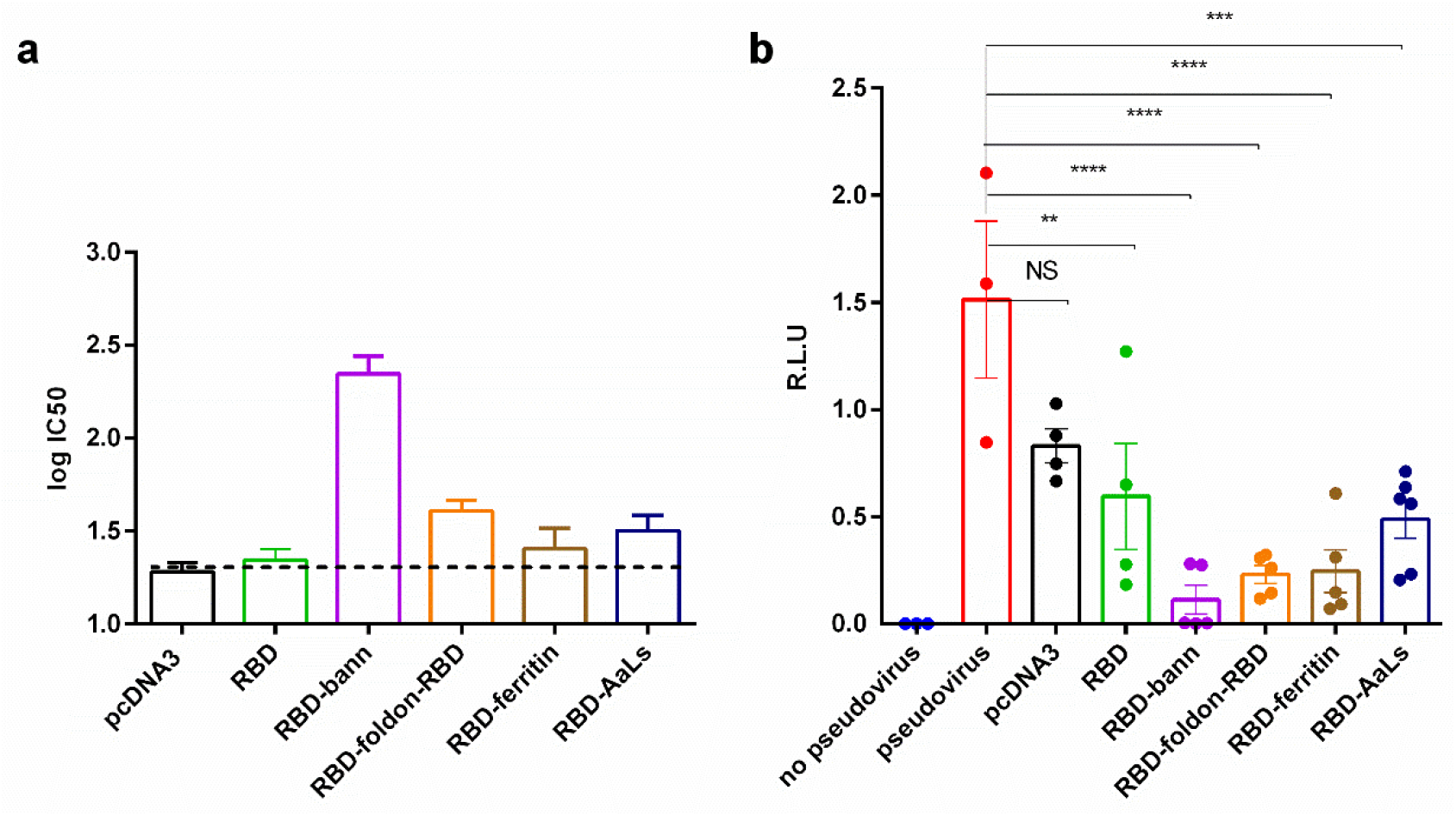
Neutralization of binding of viral RBD to the ACE2 receptor and inhibition of pseudoviral infection of cells by mouse antisera. A) Sera of mice immunized with DNA vaccines comprising scaffolded RBD were diluted and pre-incubated with Spike protein. Afterwards, Spike that bound to ACE2 was detected using streptactin-HRP. Mean and SEM of 6 (RBD-AaLs) or 5 (all others) biological replicates are shown (A) Sera of mice immunized with DNA vaccines comprising scaffolded RBD were diluted 50-fold and Spike-pseudotyped virus infection of ACE2 and TMPRSS2 –transfected HEK293 cells was followed by luminescence. Mean and SEM of 6 (RBD-AaLs) or 5 (RBD-bann, RBD-foldon-RBD, RBD-ferritin) or 4 (empty pcDNA3 vector, RBD) biological replicates are shown (B). *P< 0.05, ***P < 0.001 ****P < 0.0001. All P values are from one-way ANOVA followed by Tukey’s multiple comparisons test.

### Immunization by RBD-bann facilitates protection in the surrogate animal infection model

Mouse ACE2 is not able to bind RBD of the SARS-CoV-2, thus human ACE2 needs to be introduced to make mice sensitive to SARS-CoV-2 infection. A previous study introduced human ACE2 gene by the AAV delivery^74^. Here, we developed a model where hACE2 and TMPRRS was introduced into mice by intranasal transfection. SARS-CoV-2 Spike pseudotyped virus as used in the cell assay was used to test infection of mice in naïve as well as in immunized mice. Strong protection of pseudoviral infection in mice immunized by RBD-bann and lower in mice immunized by a monomeric RBD was observed (Fig. 6). Although this assay has several limitations as the pseudotyped virus is not able to replicate, it is nevertheless able to recapitulate the neutralization efficiency by the immunization since the antibodies target the viral recognition of the cellular receptor and, importantly, it does not require biosafety level 3 animal facilities, making it safer and faster to use by a wider range of researchers.

**Figure 6:**
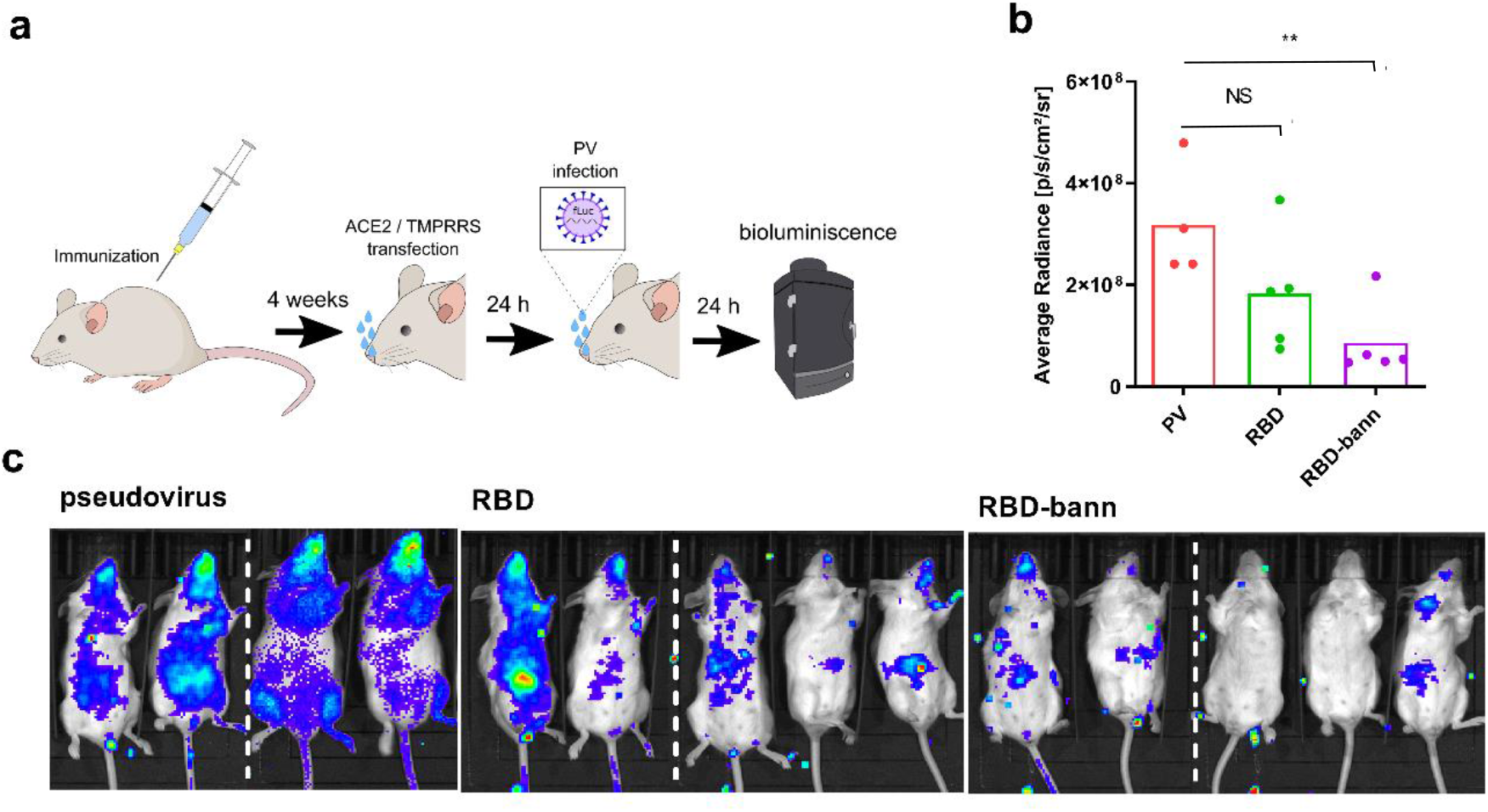
Protection of pseudoviral infection by DNA plasmid immunization in a mouse model. Mice were immunized by two injections of plasmids separated by two weeks. After one month hACE2 and TMPRRS was introduced by intranasal plasmid transfection followed by intranasal infection with SARS_CoV-2 S-typed virus (PV). Luminescence based on pseudovirus intranasal infection was measured after 24 hrs (A). Bioluminescence imaging revealing the protective state of immunized animals against pseudovirus infection in animals. Subsequent quantification of bioluminescence average radiance was carried out (B, C). Dashed line represent merging of pictures of mice from the same test group taken separately. Each dot represents an individual animal (pcDNA3 n=4; RBD and RBD-bann n=5). **P< 0.01. All P values are from Mann-Whitney test.

### Spike protein-specific cytotoxic T cells from immunized animals

An important branch of the immune response is cellular immunity mediated by T lymphocytes. Since the RBD also contains several T cell epitopes it is expected to trigger both CD4 helper as well as CD8 cytotoxic specific T cell response. After the DNA immunization we harvested mice spleens and subsequently isolated CD8 cells. By coculturing them with hACE2 transfected mouse NIH-3T3 cells that were infected with pseudovirus we observed strong specific augmented cell killing primarily for RBD-bann construct and production of γIFN (Fig. 7, Supp.Fig.3) suggesting that this type of vaccine may be able to offer protection through elimination of virus-infected cells. Also, stimulation of mouse splenocytes from DNA immunized mice, with Spike peptide pool led to highest T cell cytokine production in RBD-bann vaccinated mice (Fig. 7c)

**Figure 7:**
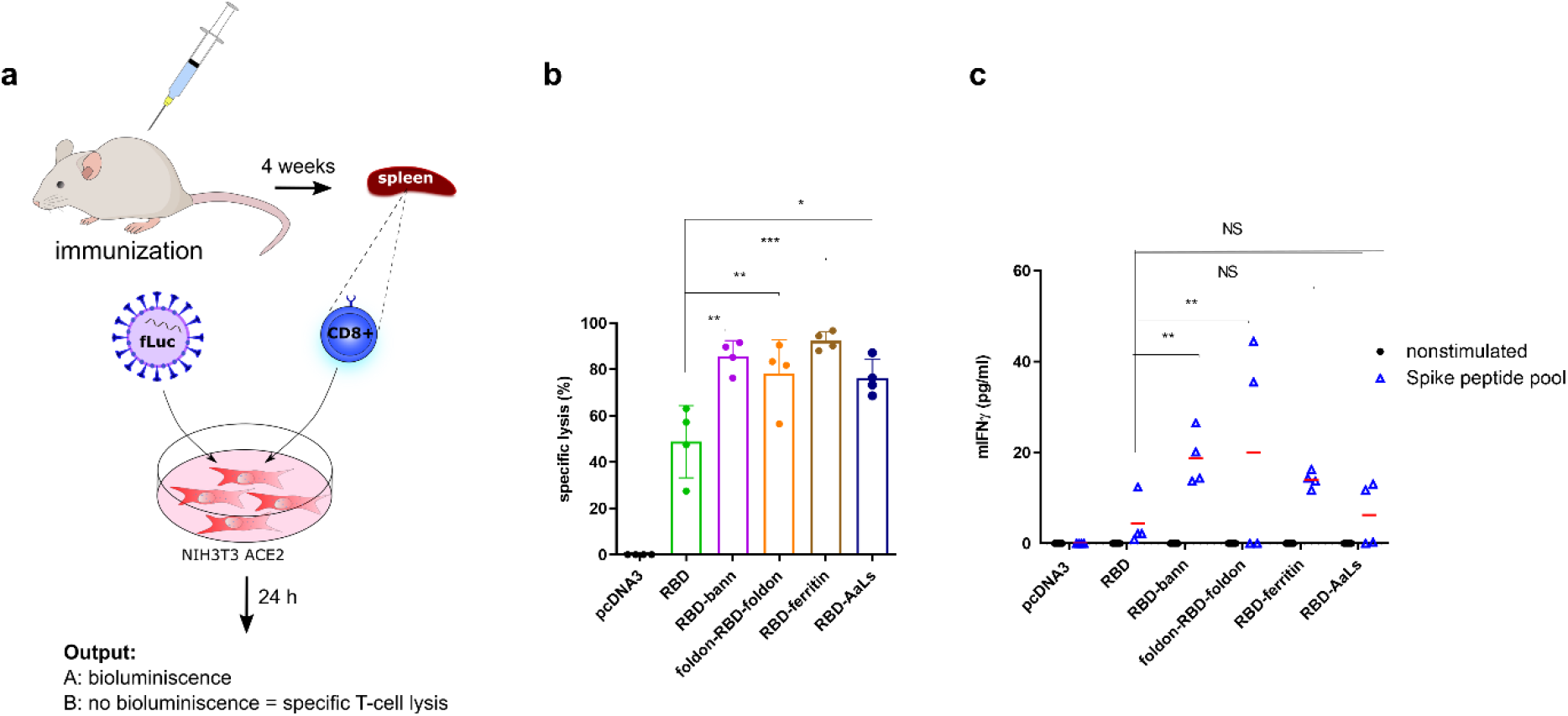
Spike protein-specific cytotoxic killing by lymphocytes from immunized animals. Cytotoxic T cell killing from immunized mice against cells expressing viral S protein. Mice were immunized with different combinations of RBD plasmid DNA and at the end of the experiment spleens were harvested. The cytotoxic effect of isolated CD8+ T cells was determined on hACE2 and TMPRRS transfected NIH-3T3, followed by pseudovirus infection. Twenty-four hours, bioluminescence was determined (A). Based on radiance values, obtained from pseuodovirus infection, specific lysis of NIH-3T3 cells was calculated (B). Mouse splenocytes from DNA immunized animals were stimulated with Spike peptide pool and 24 hours afterwards IFNγ was measured (C). Each dot represents spleen cells from an individual animal (n=4). **P< 0.01, ***P< 0.001. All P values are from one-way ANOVA followed by Tukey’s multiple comparisons test.

### Immune response against the scaffold in a genetic fusion with the target antigen

Despite many reports of different types of scaffolds for VLPs, immune response against the scaffold has been seldomly evaluated. While no effect has been reported in some cases, there are reports on the impaired immune response in case of using the same scaffolding or delivery platform^47,48^. Preexisting antibodies could opsonize the active agent in the subsequent immunization based on fusion with the same scaffold or vector and in case of genetically encoded vaccines, such as DNA, RNA or different vectors cells that produce the scaffold might be targeted by the existing cytotoxic lymphocytes. Even if it is not clear to what extent the antibodies against the scaffold hinder immune response in subsequent immunization, it should be advantageous for the scaffold to be as small as possible and hypoimmunogenic. In case of β-annulus peptide and foldon, the smallest scaffolding domains tested in this study, the response of antibodies to the immunization by the scaffolding peptides was not detectable, while lumazine-synthase exhibited strong immunogenic properties (Fig. 8). In case of foldon this is most likely due to its small size, as it has been reported to be quite immunogenic when fused to a larger cargo protein^61^. On the other hand weak response against the β-annulus peptide, which by itself forms large particles is most likely due to its hypoimmunogenicity based on its amino acid composition and therefore represents a highly suitable scaffold.

**Figure 8:**
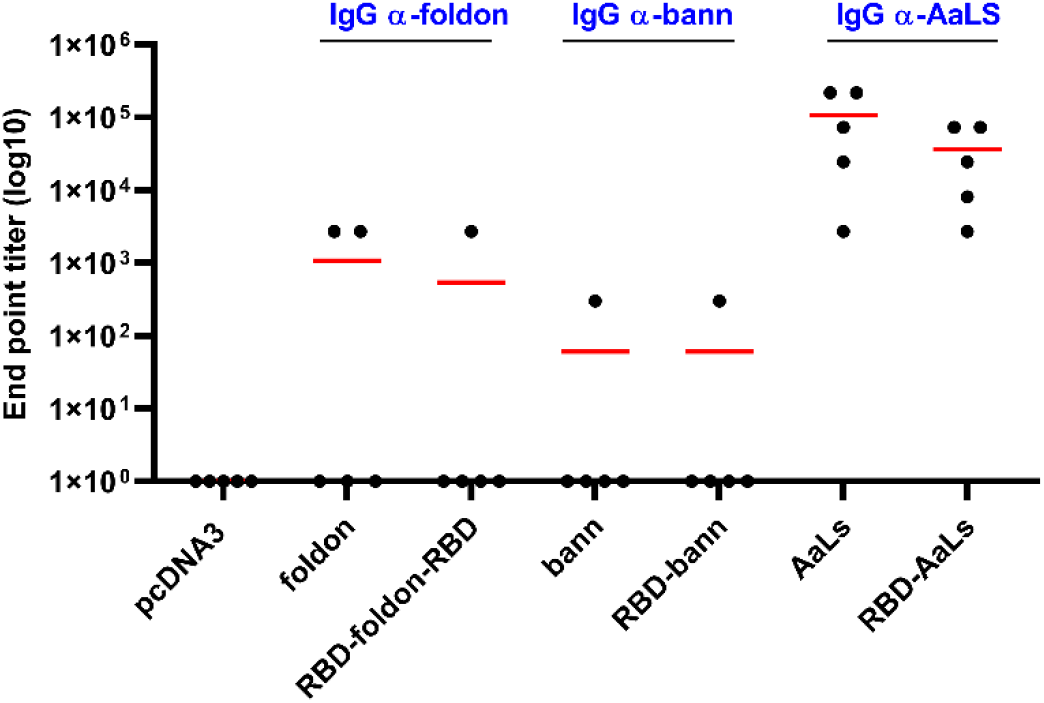
Immunogenicity of the scaffolding domains. Mice were immunized with different combination of RBD plasmid DNA. Titers of total IgG antibodies against foldon, β-annulus and lumazine synthase scaffold were tested. Graphs represent mean of EPT of group of mice (n=5 per group). Each dot represents an individual animal.

We aimed to further decrease the immune response against the scaffold by priming and booster injection of a plasmid with a different scaffold a each stage (β-annulus and foldon). Results demonstrated that the titer against the RBD remained as high as the immunization with the same scaffolded RBD (Supp. Fig. 5), while the response against the scaffold was minimal (Supp. Fig. 4).

## Discussion

Presentation of antigens, particularly their multimeric state, plays an important role in protective immune response triggered by vaccines^30,75^. Efficient presentation of antigen protein domain could improve the immune response for vaccines, such as against SARS-CoV-2, where the structural information is available on the cellular recognition. Several scaffolds for protein oligomerization with well-defined structure have been used for vaccines against different antigens, including many from viruses^31,38,39,46,65,76^. Several recent reports on vaccines using RBD exhibited high neutralization and protection against the infection^7,25^. Although even monomeric RBD protein produced a response^7^, it was substantially stronger against the dimeric RBD^25^. Here we provide a comparison of four different types of genetically encoded scaffolds fused to the RBD and implemented as DNA vaccines. We demonstrate that strongly increased antibody responses are generated already by the hexamerization scaffold, however most potent responses were obtained by scaffolds that form large oligomers. Antibody titers in this study were comparable to several other vaccines that advced to clinical trials^4,5,7,9^.The size and the degree of multimerization was not the only important factor for the observed level of neutralization as the RBD-AaLs that presents 60 copies of the antigen was less efficient in comparison to the 24 copies of the RBD presenting RBD-ferritin and 6 copies in the RBD-foldon-RBD. One reasons for the difference could be that the lumazine synthase presents the antigen in a different spatial arrangement in comparison to foldon and ferritin, while in RBD-bann assembly the presented oligomeric unit of the RBD may be governed by the intrinsic oligomerization propensity of the RBD. Scaffolded presentation of antigen generated robust T cell response which supported generation of neutralizing antibodies and generation of cytotoxic lymphocytes that could counteract the infection. This indicates that the RBD domain is sufficient as the source of T cell epitopes.

Significant attention has been devoted to the presentation of antigens on different scaffolds in a precisely defined geometry^45,77^, however the results of fusion with β-annulus peptide, that apparently generates polydisperse but large oligomers, suggest that precise positioning may not be required, unless for the generation of desired natural-like arrangement of the epitopes, such as e.g. trimerization of viral protein domains. This advantage could be explained by the wide distribution of distances between the B-cell receptors in the clusters in the fluid cellular membrane^29^. Results using DNA scaffolds to investigate the role of spatial arrangement and stoichiometry suggest that the BCR activation increases for the separation distance of the epitopes from 7-28 nm^45^. Even in our smallest hexameric assembly the distance between the epitopes was around 8 nm, and exceeded 27 nm in larger assemblies (e.g. RBD-AaLS). Insoluble aggregates based on amyloid fibril-forming peptides as the scaffolds for vaccines^50,53^ are likely not efficiently transported to the lymph nodes and germinal centres^34,78^. An additional explanation for the efficiency of the β-annulus scaffolded vaccine may be a dynamic equilibrium between the oligomeric states of the assembly, which may facilitate the trafficking and presentation to B-lymphocytes. T cell response as shown here also plays an important role in both generation of antibodies as well as for the cytotoxic activity.

Small scaffolds have the advantage of maximizing the fraction of the antigen and are more likely to be hypoimmunogenic, avoiding nonproductive or potentially even harmful diversion of the immune response. While foldon has been reported to be strongly immunogenic in the context of fusion to other proteins and attempts were made to silence its immunogenicity by glycosylation^61^, its positioning at the core of the RBD-foldon-RBD assembly likely decreased its exposure to the B-cell receptors. The other tested small scaffold, β-annulus peptide contains predominantly weakly immunogenic residues and our results demonstrate that immunization with β-annulus peptide that forms large assemblies resulted in a weak immune response. Vaccination with AaLs-scaffolded RBD on the other hand generated strong antibody response against the scaffold.

DNA vaccines have demonstrated particularly good efficiency as priming agents in protein booster combination^79,80^, which could be used as an implementation for the vaccine presented. Combinations of vaccines based on different scaffolds for the prime and boost step could additionally decrease the immune response against the target and focus the response to the target RBD domain. Further, prime-boost combinations that involve DNA vaccine have been shown to boost mucosal immunity^80–82^ suggesting therapeutic advantage for Covid-19.

While we have used DNA delivery platform to induce production of oligomeric RBD in situ in the tissue, the same scaffolding strategy and domains could be used for other delivery platforms, including isolated proteins, mRNA as well as diverse viral platforms as the scaffolding improved presentation of antigens and immune response. Nevertheless it needs to be investigated if this strategy of presentation is efficient also for other protein antigens.

As a conclusion genetic fusion of target antigen to small scaffolding domain enables fast design and implementation as nucleic acid based vaccines particularly in case of epidemic emergencies. Assemblies that present multiple copies of the RBD as presented here are promising candidates for clinical studies as a vaccine against Covid19 and represent a platform for vaccines against other emerging viral diseases.

## Acknowledgments

This work was supported by the Slovenian Research Agency (program P4-0176, projects Z3-9276, Z3-9260), H2020 project Virofight and ERC AdG project MaCChines. We would like to thank Markus Hoffmann for his advices on pseudoviral system. We are grateful to Stefan Pölhmann for pseudoviral system and plasmids. We would like to thank Tina Šket, Eva Koplan, Neža Pavko, Veronika Mikolič, Jure Bohinc, Irena Škraba and Anja Perčič from NIC for their technical support and Laurence Durrer and Soraya Quinche from PTPSP-EPFL for production in mammalian cells.

---

**Supplementary Figure 1:**
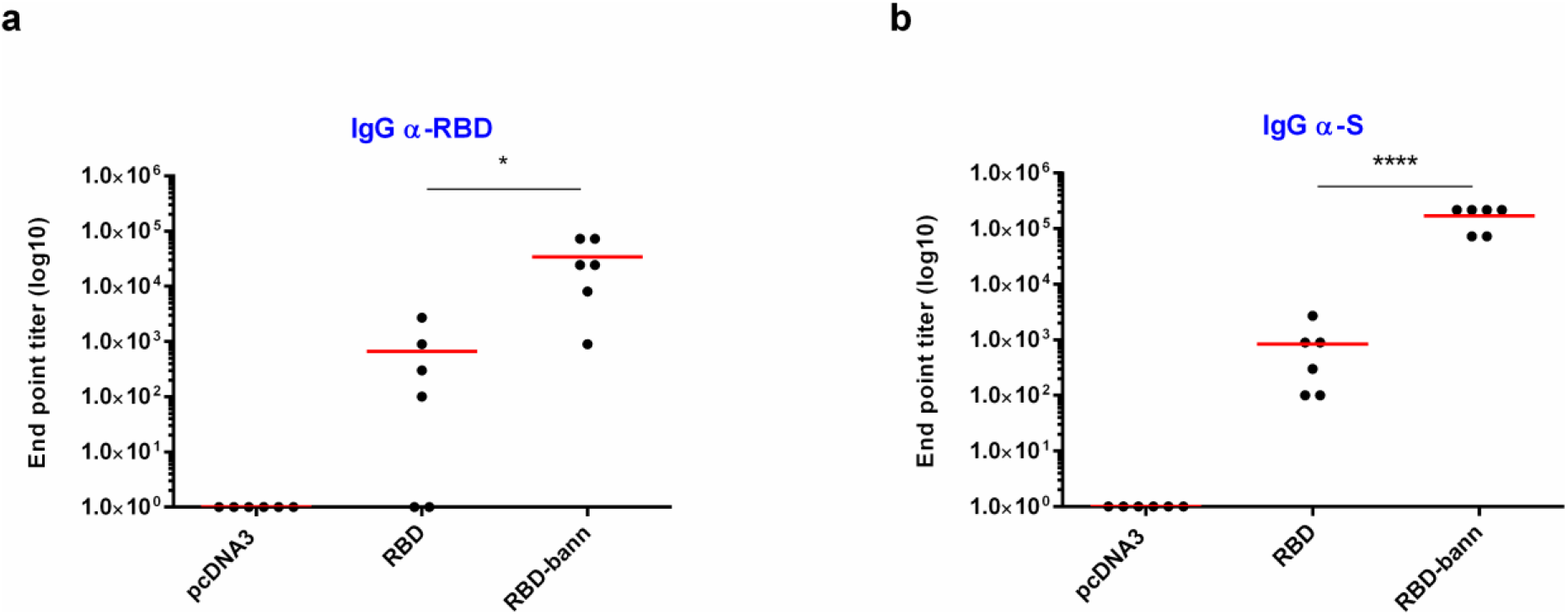
DNA plasmid immunization with naked DNA. Mice were immunized with 20 μg per animal of naked DNA (empty vector, RBD, RBD-bann), dissolved in 150 mM NaCl. End point titer (EPT) for total IgG against RBD (A) and against Spike protein (B) were determined by ELISA. Graphs represent mean of EPT of group of mice (n=6 per group). Each dot represents an individual animal. *P< 0.05; ****P < 0.0001. All P values are from one-way ANOVA followed by Dunnett’s multiple comparisons test.

**Supplementary Figure 2:**
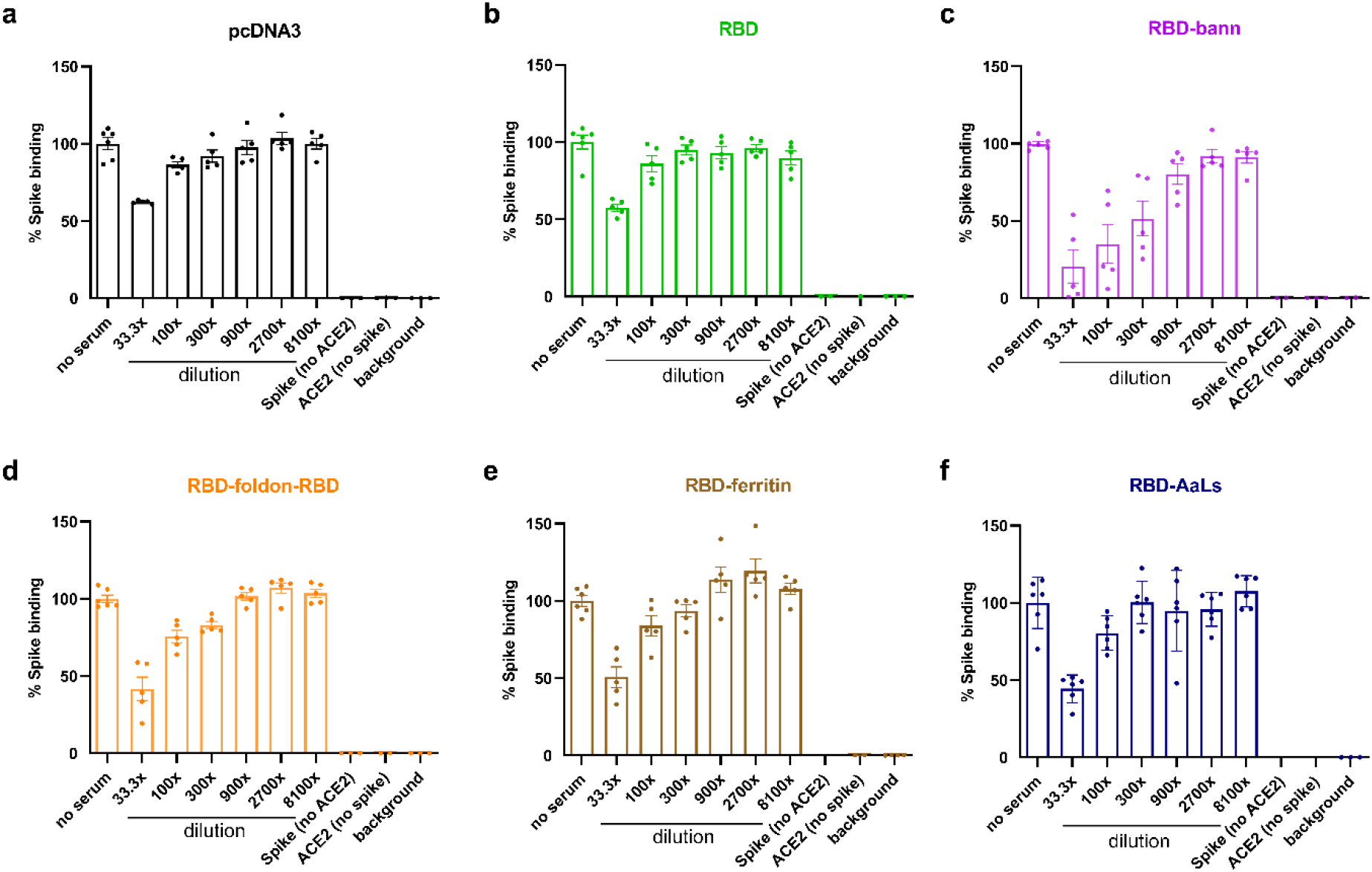
Neutralization assay based on inhibition of ACE2 Spike interaction in the presence of sera of DNA immunized mice. Different mice sera dilutions were incubated with Spike protein. By ELISA test, inhibition of Spike and ACE2 protein was determined, indicating neutralization effect of anti-Spike antibodies, found in DNA immunized mice.

**Supplementary Figure 3:**
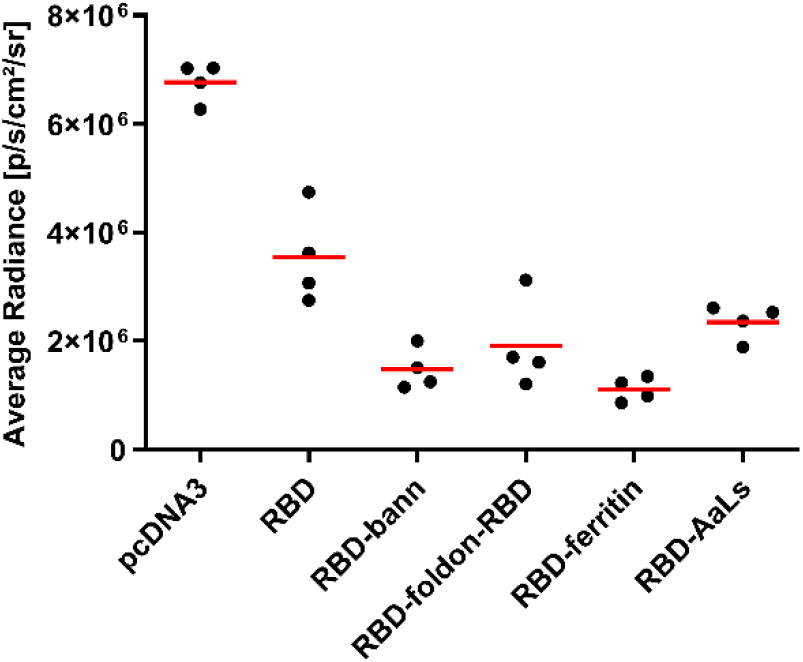
Bioluminscence measurement of pseuodvirus infection of hACE2 and TMPRRS transfected NIH-3T3 after the addition of mouse isolated CD8+ cells. Twenty-four hours later bioluminscece was measured in cell coculture of mouse isolated CD8+ cells and NIH-3T3 cells, infected with pseudovirus.

**Supplementary Figure 4:**
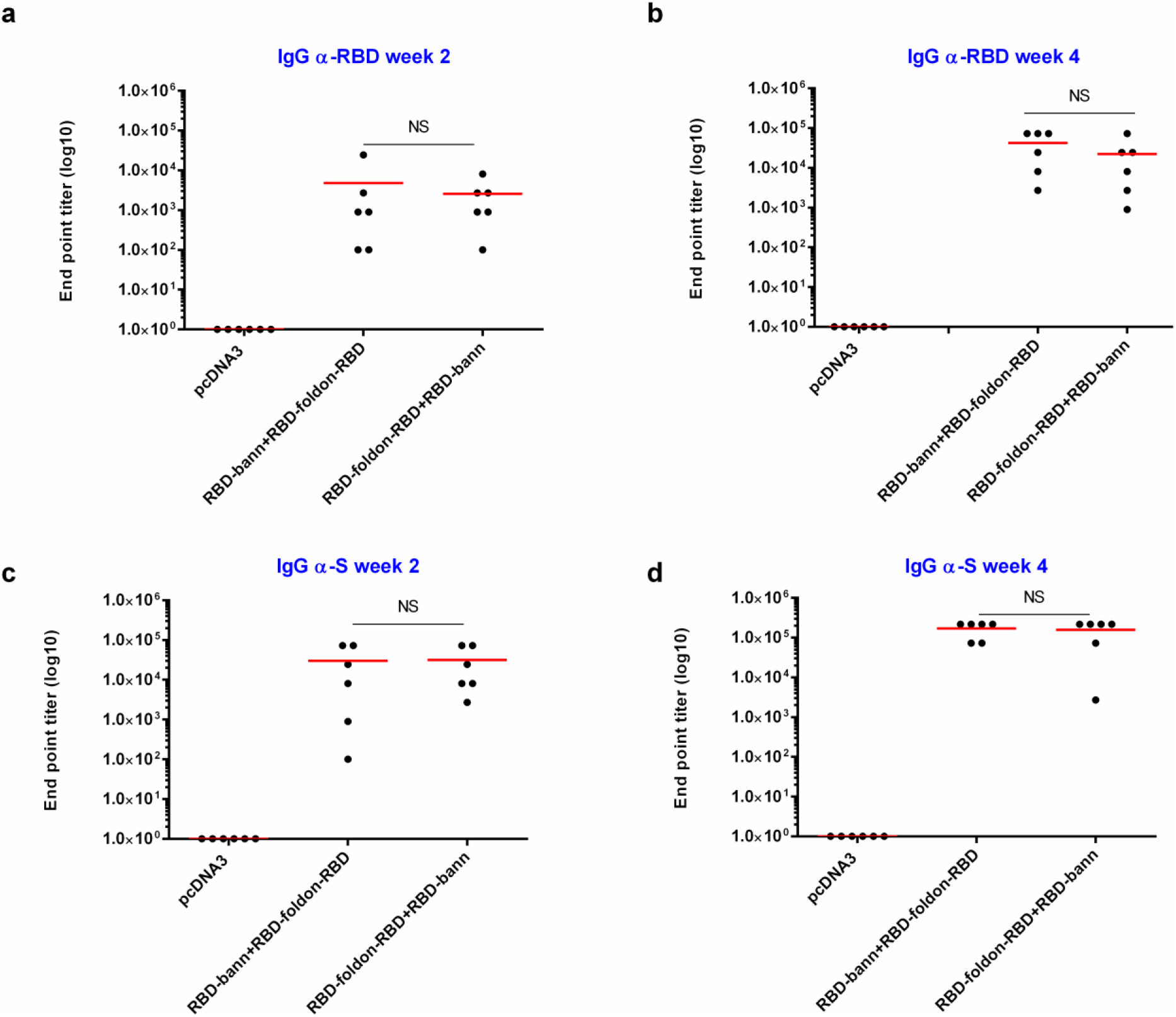
Total IgG in mice that underwent switch immunization. Mice were immunized with combinations of differently scaffolded RBD plasmid DNA (β-annulus and foldon) for prime and boost immunization. Titers of antibodies against RBD after prime and boost (A,B) and against Spike protein (C, D) were determined via ELISA. Graphs represent mean of EPT of group of mice (n=6 per group). Each dot represents an individual animal. To determine NS, Mann-Whitney test was performed.

**Supplementary Figure 5:**
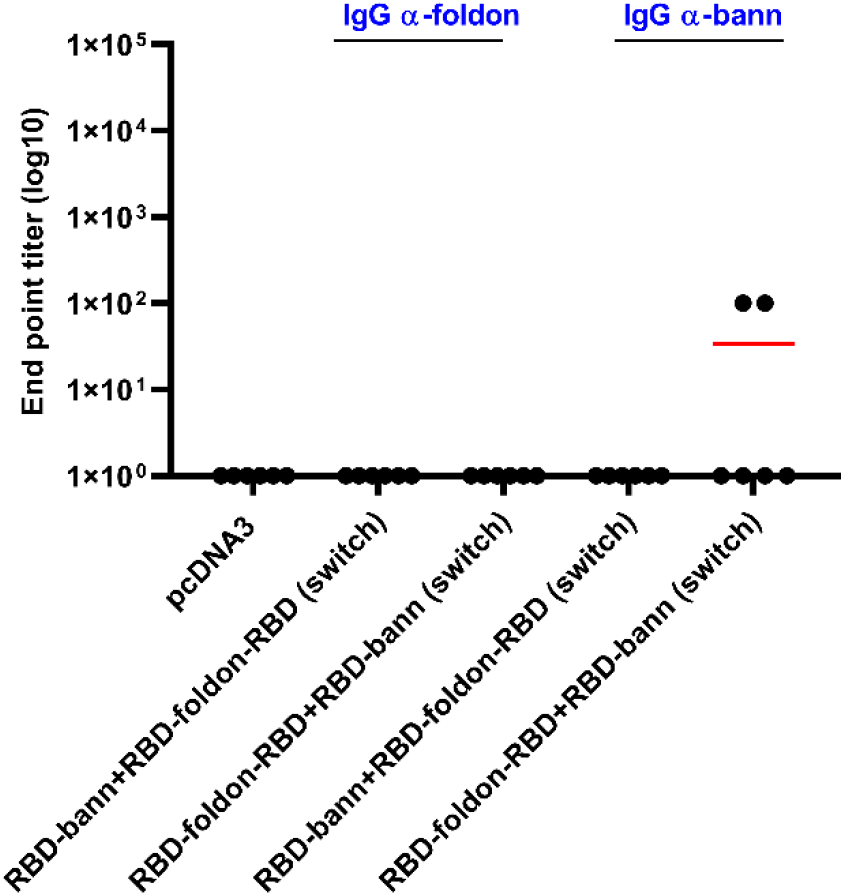
Total IgG against scaffold in mice that underwent switch immunization. Mice were immunized with combinations of differently scaffolded RBD plasmid DNA (β-annulus and foldon) for prime and boost immunization and vice versa. Titers of antibodies against scaffold (depicted in blue) after prime and boost were determined via ELISA. Graphs represent mean of EPT of group of mice (n=6 per group). Each dot represents an individual animal.

